# Gravitational and dynamic components of muscle torque underlie tonic and phasic muscle activity during goal-directed reaching

**DOI:** 10.1101/138263

**Authors:** Erienne Olesh, Bradley Pollard, Valeriya Gritsenko

## Abstract

Human reaching movements require complex muscle activations to produce the forces necessary to move the limb in a controlled manner. How gravity and the complex kinetic properties of the limb contribute to the generation of the muscle activation pattern by the central nervous system (CNS) is a long-standing question in neuroscience. To address this question, muscle activity is often subdivided into static and phasic components. The former is thought to be related to posture maintenance and transitions between postures. The latter represents the remainder of muscle activity and is thought to be related to active movement production and the compensation for the kinetic properties of the limb. In the present study, we directly addressed how this subdivision of muscle activity into static and phasic components is related to the corresponding components of active muscle torques. Eight healthy subjects pointed in virtual reality to visual targets arranged to create a standard center-out reaching task in three dimensions. Muscle activity and motion capture data were synchronously collected during the movements. The motion capture data were used to calculate gravitational and dynamic components of active muscle torques using a dynamic model of the arm with 5 degrees of freedom. Principal Component Analysis (PCA) was then applied to muscle activity and the torque components, separately, to reduce the dimensionality of the data. Muscle activity was also reconstructed from gravitational and dynamic torque components. Results show that the gravitational and dynamic components of muscle torque represent a significant amount of variance in muscle activity. This method could be used to identify static and phasic components of muscle activity using muscle torques. The contribution of both components to the overall muscle activity was largely equal, unlike their relative contribution to active muscle torques, which may reflect a neural control strategy.

## Introduction

The musculoskeletal anatomy of the body constitutes a complex dynamical system that is a challenge to control for the central nervous system (CNS). Some of the complexity is due to the muscular redundancies that allow humans to perform complex tasks. Additional complexity results from the forces associated with the mechanical properties of the multi-joint limb, termed limb dynamics. How the CNS deals with limb dynamics is commonly investigated through joint torques, or rotational forces, that arise during motion of the limb (Flanders, 1991; Sainburg et al., 1995; 1999; Shabbott and Sainburg, 2008) or from action of external forces when the limb is held stationary (Buneo et al., 1997; Weiss and Flanders, 2004). During these tasks, angular kinematics (position and velocity) can be used to derive joint torques for each independent direction of motion termed degree of freedom (DOF) using equations of motion. The goal is to derive the active torques that are the result of muscle contractions in the presence of passive forces that have both extrinsic and intrinsic sources (Dounskaia and Wang, 2014; Gentili et al., 2007; Le Seac’h and McIntyre, 2007; Papaxanthis et al., 2005). A large contributor to the extrinsic passive torques is gravity. These gravitational torques depend on the orientation of limb segments in space, and thus they contribute to both posture and movement (Bastian et al., 1996). The compensation for gravitational torques is important for motor control, as evidenced by altered patterns of movement errors and muscle activity of people moving in micro-gravity environments (Fisk et al., 1993; Papaxanthis et al., 1998; 2005; Pozzo et al., 1998). Gravity torques can also be optimally integrated in the planning of rapid arm movements and exploited to reduce muscular efforts during rapid motions (Gaveau et al., 2014; 2016; Rousseau et al., 2016). Studies of cerebella pathologies and adaptation after returning from microgravity environment to normal gravity have also suggested that the effect of gravity on the arm may be separately estimated from the effect of dynamic torques (Gaveau et al., 2011; Sajdel-Sulkowska, 2013). The proportion of active muscle torques that is responsible for gravity compensation can be estimated as the difference between muscle torques produced in a micro-gravity environment and muscle torques produced under normal gravity. This portion of muscle torques has a different temporal profile than that of the motion-related dynamic components of muscle torque (Russo et al., 2014). In another definition, the dynamic component of muscle toque varies with the speed of movement, while the gravitational component of muscle torque does not (Flanders and Herrmann, 1992; Hollerbach and Flash, 1982).

Traditionally, the postural transition component of muscle activity has been estimated as a linear ramp in electromyography (EMG) during movement between EMG values obtained before and after movement, i.e. during posture maintenance (Buneo et al., 1994; Flanders et al., 1996). This static component is often subtracted from the EMG during movement, and the residual phasic EMG is studied as the motion-related signal. While the estimate of postural EMG are valid, there is no physiological evidence for a linearly-changing EMG associated with the transition between postures during movement. An improvement on this technique would be a quantitative estimate of the contribution of gravity acting on the limb during movement to muscle activity. In addition, relating phasic EMG to the dynamic component of muscle torque would be useful in evaluating the contribution of individual muscles to active torques responsible for movement vs. joint stiffness, that is not accounted for by active torques. Damage to the cerebellum appears to uniquely affect the phasic component of EMG in a way that supports its role in controlling passive torques (Manto and Bosse, 2003). In the current study, we examine the contribution of gravitational and dynamic components of muscle torques to EMG during goal-directed reaching movements. EMG of different muscles and torques about different DOFs are coupled through the kinematic chain of the limb. We control for this coupling by reducing the dimensionality of our data using principal component analysis (PCA). We use PCA to obtain independent components from muscle torques and compare them to the independent components obtained from EMG, as well as calculate the amount not variance torque components account for in EMG. We expect that gravitational and dynamic torque components capture significant amounts of variance in EMG and their waveforms can be used to identify static and phasic components in EMG.

## Methods

Eight healthy individuals (5 males, 3 females) with an average age of 24.8 ± 0.71 years old were recruited to perform a reaching “center-out” task. The study and the consent procedure were approved by the Institutional Review Board of West Virginia University (Protocol # 1311129283). All subjects provided their written consent prior to participating in the study. All subjects were right-hand dominant and reported no movement disorders and no major injuries to their right arm. Height, weight, and arm segment lengths were measured for each subject and used to adjust model parameters to create subject-specific dynamic models (see below).

Movements were instrumented using a virtual reality software (Vizard by Wolrdviz) and head set (Oculus Rift), which displayed 14 targets arranged in two perpendicular planes, horizontal transverse plane and vertical coronal plane (Fig 1). To reduce inter-subject variability in kinematic data, the target locations were adjusted for each subject based on the lengths of their arm segments, which ensured the same initial and final joint angles across all movement directions. The center target was placed so that initial arm posture was at 0-degree shoulder flexion, 90-degree elbow flexion, and a 0-degree wrist flexion. The distance from the center target to the peripheral targets was scaled to 30 percent of each subject’s total arm length (from anterior acromial point to the distal end of the index finger). On average, this amounted to 20 cm distance from the central to peripheral targets. This scaling reduced the inter-subject variability in the joint angles at each peripheral target. Each movement began with the subject pointing to the center target, which was the only one visible. After VR detected the tip of the subject’s finger inside the target radius, the central target changed color and one peripheral target appeared. When the VR detected the tip of the subject’s finer inside the peripheral target radius, it changed color, which cued the subject to return to the central target. Upon returning to the central target the task reset, peripheral target disappeared and a new one appeared after a delay of 0.5 s. Subjects were instructed to not move the trunk, keep their wrist pronated and straight, and point as quickly and accurately as possible. Movements to each target location were repeated 15 times and performed in a randomized order.

**Figure 1.**
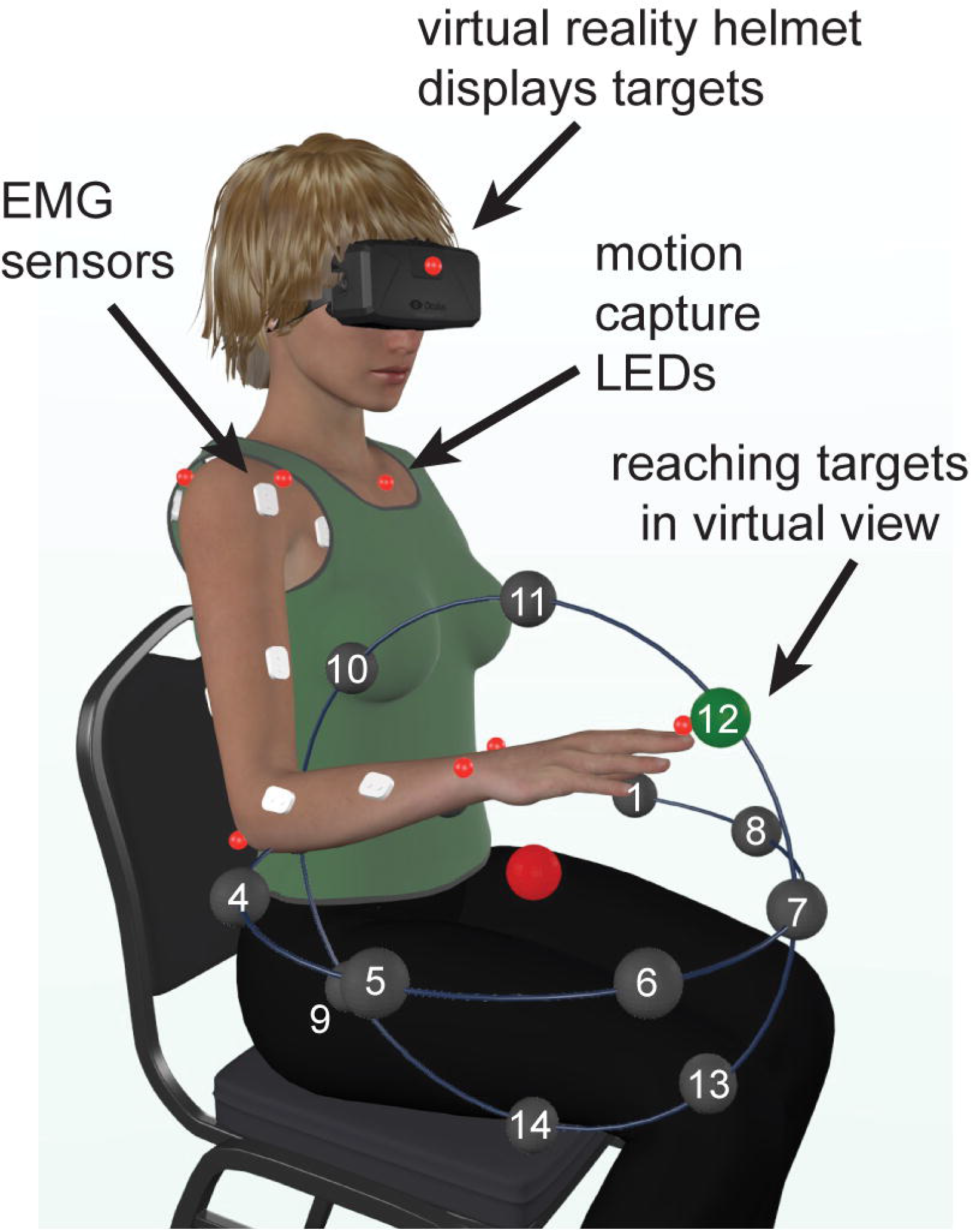
Experimental setup. Illustration showing the locations of reaching targets, arranged in a semi-spherical pattern in VR, relative to the physical location of the subject. The central target is shown in red and one of the goal targets is shown in green.

Arm and trunk movements were recorded with an active motion capture system (PhaseSpace, Impulse) at 480 frames per second. The light emitting diodes of the motion capture system were placed on anatomical landmarks according to best practice guidelines {Robertson:2004wn}. Electromyography (EMG) was recorded from twelve arm muscles at a rate of 2000 Hz (MA400-28 MotionLab Systems). Muscles recorded during the experiment included the pectoralis major (Pec), teres major (TrM), anterior deltoid (AD), posterior deltoid (PD), long and short heads of the biceps (BiL and BiS respectively), lateral and long heads of the triceps (TrLa and TrLo respectively), brachioradialis (Br), flexor carpi ulnaris (FCU), flexor carpi radialis (FCR), and extensor carpi radialis (ECR). Motion capture and electromyography were synchronized using a custom circuit and triggering mechanism (Talkington et al., 2015). Motion capture and EMG data were imported into Matlab and processed as follows using custom scripts.

Digitized EMG data were high pass filtered at 20 Hz to remove motion artifacts, rectified, and low pass filtered at 10 Hz, consistent with SENIAM recommendations. Motion capture data were low pass filtered at 10 Hz and interpolated with a cubic-spline. The maximum interpolated gap was 0.2 seconds. The onset and offset of movement was found based on the velocity of three hand LEDs changing by five percent of the maximum velocity for a given movement. These events were used for temporal normalization of all data. Signals starting 200 ms prior to the onset of movement were included in all analyses to ensure adequate capture of initial EMG bursts and onset of phase-advanced torques. Arm kinematics were obtained from motion capture by calculating Euler angles and angular velocity for five joint DOFs including shoulder (flexion/extension, abduction/adduction, pronation/supination), elbow (flexion/extension), and wrist (flexion/extension). Hand pronation/supination and wrist abduction/adduction were found to be minimal during the pointing task, and thus these DOFs were not included in the analysis.

## Limb dynamics

To calculate joint torques, an inverse dynamic model of the subject’s arm was constructed in Simulink (MathWorks). The model comprised 5 DOFs as described above and three segments approximating inertial properties of the arm, forearm, and hand. Motion of the trunk was found to be minimal during the task, thus the model had the trunk fixed in space. Inertia of the segments was approximated with a cylinder of the length equal to that of the corresponding segment and a 3 cm radius. The masses and centers of mass for each segment were determined by their anthropometric ratios to the subjects’ segment lengths and weight (Winter, 2009). The model implemented equations of motion that can be summarized as follows:

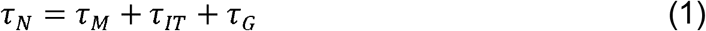

, where τ_N_ is a vector of net torques that produce movement; τ_M_ is a vector of active torques due to muscle contractions; τ_IT_ is a vector of passive interaction torques; τ_G_ is a vector of passive torques caused by gravity.

Angular kinematics averaged per movement direction and per subject was used in the subject-specific inverse model to calculate muscle torques (Fig. 1B), similar to that in (Russo et al., 2014). This is equivalent to rearranging the equation (1) as follows:

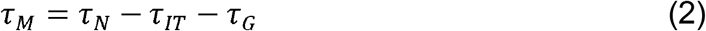

The computed muscle torques are proportional to the sum of all moments generated by muscles spanning the joints:

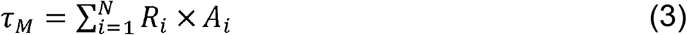

, where *R_i_* is moment arm of muscle *i* about a given DOF, and *A_i_* is activation of muscle *i*. The numerical quality of inverse dynamic simulations was checked by running the same model in forward dynamics mode using the calculated torques as inputs and simulated angular kinematics as outputs. The simulated and experimental joint kinematics was compared, and the mean ± standard deviation of the root-mean-squared differences between them was 0.05 ± 0.02 radians across all DOFs.

As described in the Introduction, muscle activity is often separated into static and phasic components. Here we hypothesized that gravitational and dynamic components of muscle torque may underlay the static and phasic components of EMG respectively. This can be represented as follows:

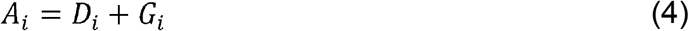

and substituting equation (4) into (3) gives

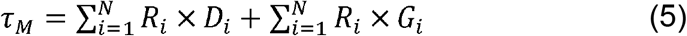

Because torques are additive, the muscle torques obtained using the inverse model can be separated into two components (Gottlieb et al., 1997; Russo et al., 2014), which could be proportional to the dynamic and gravitational components of EMG. To estimate the dynamic component of muscle torques responsible for motion production and inter-joint coordination without gravity, the inverse model was run without simulating external gravitational force (the parameter for gravitational force in the physics engine was set to 0). This resulted in the following:

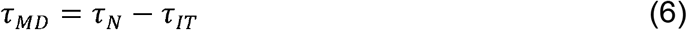

, where τ_MD_ are muscle torques that would produce the same motion without gravity as that recorded in the presence of gravity. Example of such torques would be the sum of muscle moments produced during motion in microgravity environment. Another example are torques necessary for planar movements in a horizontal plane, where the force of gravity is perpendicular to the plane of motion (Debicki and Gribble, 2005). Then the component of muscle torque that is needed to compensate for gravity (τ_MG_) can be estimated as the difference between muscle torques with and without gravity as follows:

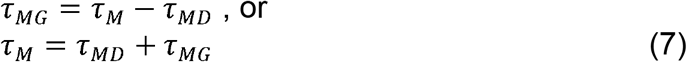

Below, τ_MD_ is referred to as MD torque, while τ_MG_ is referred to MG torque for simplicity.

The relative contribution of MG and MD torques to the overall muscle torques was calculated as the shared variance between each torque component and muscle torque for a corresponding DOF. Muscle, MG, and MD torques were averaged per movement direction per subject, and the coefficient of determination (r^2^) was used to quantify shared variance between MG and muscle torques and separately between MD and muscle torques for corresponding DOFs. These r^2^ were divided by 2 to bring them on the same scale as r^2^ calculated for EMG decomposition as described below.

## Dimensionality Reduction

To control for widespread correlations between biological signals, EMG and dynamic data were reduced in dimensionality using principal component analysis (PCA). Rectified EMG signals were normalized to movement duration, averaged per movement direction, and low pass filtered at 10 Hz. To ensure that muscle activations were unitless, maximum contraction values were calculated for each muscle across all movement directions and used to divide mean EMG for each movement direction. The resulting data matrix was comprised of 336 columns (12 EMG signals for 14 movements toward each virtual target and 14 return movements) and 1000 rows representing samples in time. To ensure that MD and MG torques were unites, the maximal amplitudes across all movement directions were used to divide torques for each movement direction. The MD and MG torque data matrices were comprised of 140 columns each (torques for 5 DOFs for 14 movements toward each virtual target and 14 return movements) and 1000 rows representing samples in time. All data were demeaned; eigenvalues and eigenvectors were obtained using singular value decomposition in Matlab. Eigenvectors were direction independent waveforms in time, while eigenvalues represented projections of signals onto the eigenvectors per muscle or DOF per movement direction.

To evaluate the contribution of gravity and dynamic torques to muscle activation, the first eigenvectors that captured the most variance in MD and MG data were used to decompose EMG data. Projections of EMG data onto the torque eigenvectors, the z-scores, were calculated using a least-squares method. The EMG data were then reconstructed back from the obtained z-scores and torque eigenvectors. The coefficient of determination (r^2^) was used to quantify the quality of EMG reconstruction. Separate r^2^ were calculated to evaluate the amount of shared variance captured by each of the torque eigenvectors. These r^2^ were divided by their sum and multiplied by the r^2^ of the overall EMG reconstruction.

## Statistical analysis

Statistics on r^2^ values was done using repeated measures analysis of variance (rANOVA) in MATLAB. A single rANOVA model was fitted to the r^2^ from EMG decomposition and the r^2^ from torque components across all signals and movement directions per subject. Only EMG signals per movement direction per subject for which the peak of mean EMG activity was larger than 30% of maximal value across movement directions were included in this analysis. This ensured that only active muscles were analyzed. The model utilized a within-subject design with 3 factors. The TM factor grouped data based on torque (PCA eigenvalues) or muscle (EMG decomposition z-scores). The DG factor grouped data based on the type of component (dynamic or gravitational). The movement factor grouped data based on the direction of movement (14 unique directions in Fig. 1A). Post-hoc multiple comparisons were used to further examine significant interactions. Linear regressions between the r^2^ from EMG decomposition and the r^2^ from torque components were used as measures of the contribution of MG and MD torques to EMG.

Data trends are reported using means ± standard deviations across subjects, which are included in the Results section below unless otherwise specified.

## Results

The motion in virtual reality was highly consistent, as demonstrated by the low standard deviations of angular kinematics across the fifteen repetitions of each movement (Fig 2A). The mean endpoint error was 0.04 ± 0.01 m across subjects. The kinematic profiles showed typical motor invariance with bell-shape velocity profiles with peaks ranging from 1.2 to 3.3 meters per second, and accompanying acceleration and deceleration phases (Fig. 2A). The movements were produced by active muscle torques, whose temporal profiles were subdivided into gravitational MG and dynamic MD components as described in the Methods (Fig 2B). The MG component waveform changed in a single direction during a given movement and was the source of the offset in the muscle torque waveforms. The waveform of the MD component was largely similar to the angular acceleration waveform. As expected, muscle activity was more variable across subjects, but most muscles did follow a reciprocal pattern of activation for movements in the opposite directions (Fig. 2C, first 2 columns vs. the last 2 columns).

**Figure 2.**
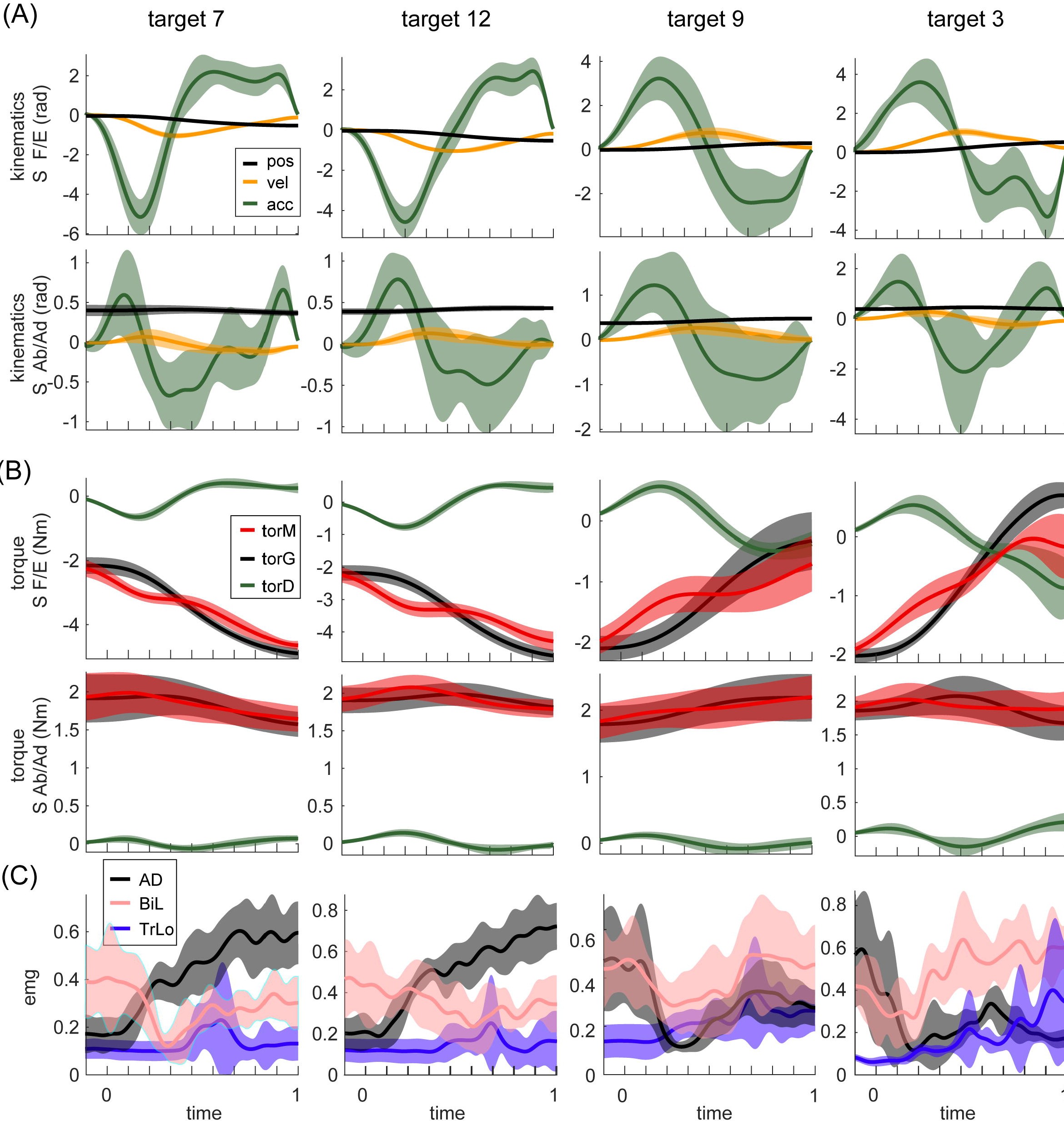
Example motion signals and muscle activity, for a single subject. Columns show signals from movements to four different targets. Targets are numbered as in Fig. 1. Movements in the first 2 columns are adjacent to each other and orthogonal to those in the other 2 columns (see Fig. 1A). Lines are averages across 15 repetition of the same movement; shaded areas are standard deviations. (A) The temporal profiles of kinematic signals for two DOFs, shoulder flexion/extension and abduction/adduction are shown. (B) Dynamic signals calculated from signals in (A). (C) EMG signals from three muscles for the corresponding movements. Muscle abbreviations are as described in the Methods section.

The linear dependencies across torque components and across EMGs were examined using PCA as described in Methods. In MG torques across all movements, a single principal component accounted for 96 ± 1 % of variance. The second principal component accounted for 4 ± 1 % variance, and the rest accounted for progressively less variance. For MD torques across all movements, a single principal component accounted for 92 ± 2 % of variance. The second principal components accounted for 6 ± 2 % of variance. In contrast, for EMG the first three principal components accounted for a comparable amount of total variance, 66 ± 22 %,11 ± 4 %, and 5 ± 2 % of variance accounted for by the principal components 1 through 3 respectively. The waveforms (eigenvectors) of the first and second principal components of EMG were very similar to the first principal components of MG and MD torques respectively (Fig. 3A). Therefore, we used the first principal components of MG and MD torques to decompose EMG data and calculate the variance accounted for by these dynamic signals.

**Figure 3.**
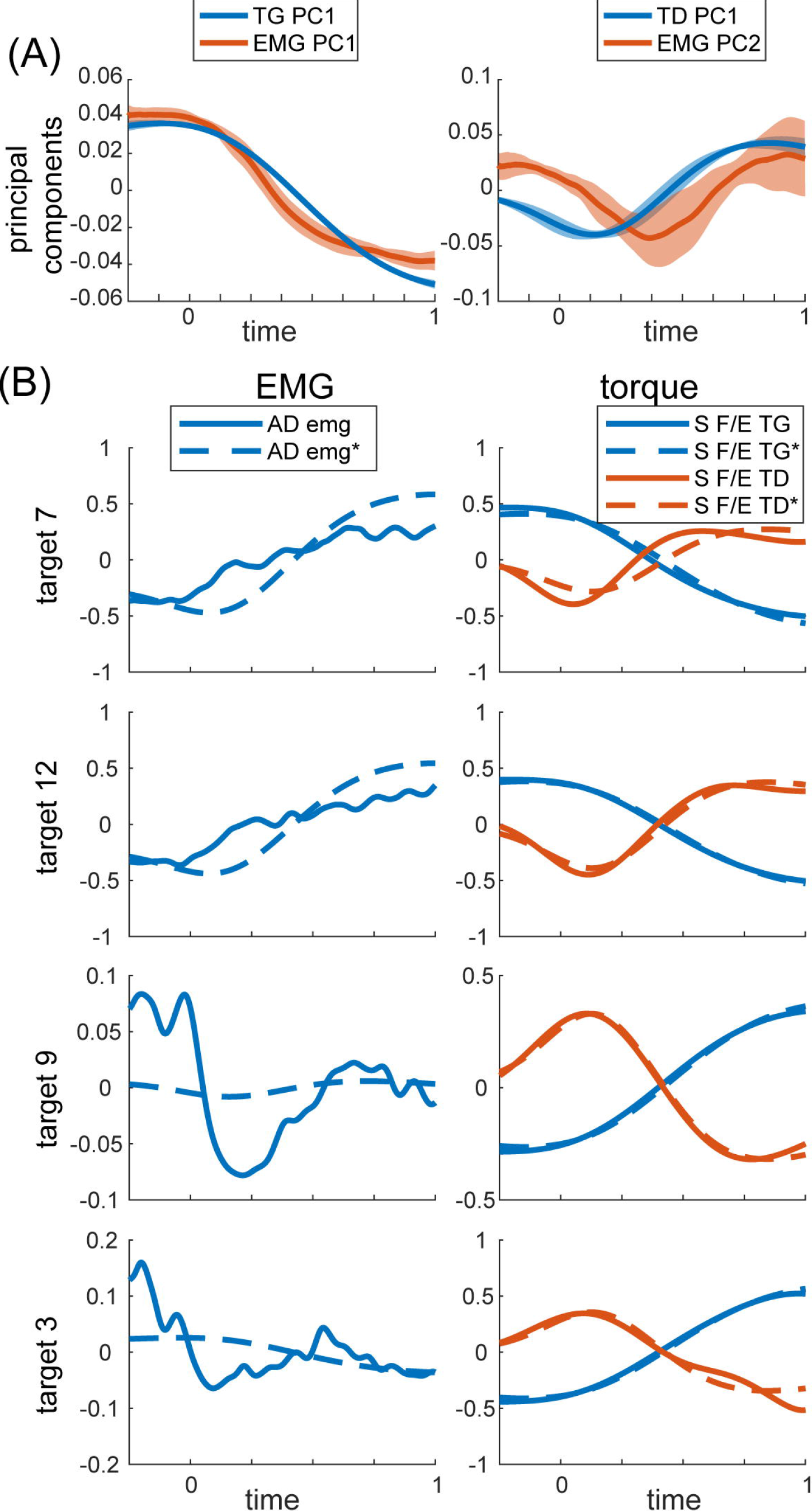
PCA on torques and EMG. (A) Temporal activation profiles of EMG and torque components. Average normalized activation profiles (solid lines) and standard deviations (shaded area) across all eight subjects are plotted for each eigenvector. (B) Example recorded and reconstructed signals from the same subject and movements as in Fig. 2. MG and MD torques were reconstructed from their respective first principal components only. EMG was reconstructed from the same torque components as described in Methods. MG and MD torques and EMG were normalized and demeaned as described in Methods. Reconstructed signals are labeled with * and marked with dashed lines. S F/E stands for Shoulder flexion/extension torque.

The total variance accounted for by torque decomposition of EMG was 61 ± 23 % across subjects. These values varied between individual movement directions (Fig. 4A). However, the relative contribution of gravitational and dynamic principal components to EMG varied independently from the relative contribution of MG and MD torques to the overall muscle torques across the different movement directions (Fig. 4B & C). The relative variance in EMG accounted for by each torque principal component was lower than the relative variance that MG and MD torques contribute to the muscle torque (rANOVA: difference % 12%; standard error % 3%; p % 0.036). This difference was due to the dynamic principal component accounting for less variance in EMG than expected from MD contribution to muscle torque (rANOVA: difference between means % 19%; standard error % 5%; p % 0.027) compared to those for gravitational components (rANOVA: difference between means % 5%; standard error % 2%; p % 0.116). There was no average difference between the relative contributions of the gravitational and dynamic components across all movement and signals (rANOVA: difference % 3%; standard error % 2%; p % 0.36). For individual muscles, the total and relative variances accounted for by torque components was largely the same (Fig. 5).

**Figure 4.**
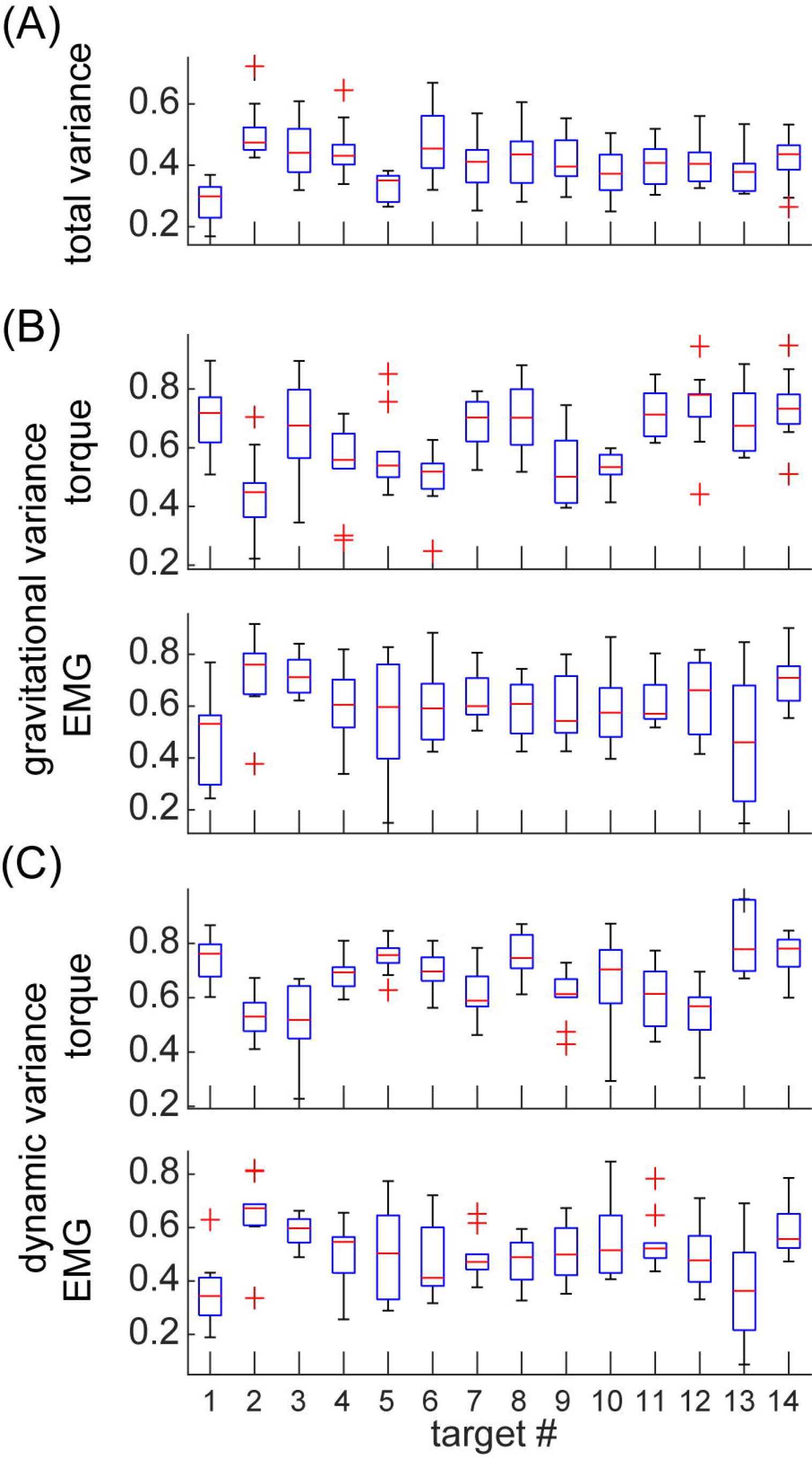
Variance accounted for by the individual components in EMG and torques per movement direction. Values are means across subjects and signals; boxes indicate ranges of 25 and 75 percentiles in data; error bars show standard deviations; red crosses indicate outliers. (A) Total variance accounted for by torque decomposition of EMG per movement direction. (B) Relative variance accounted for by gravitational principal component. (C) Relative variance accounted for by dynamic principal component.

**Figure 5.**
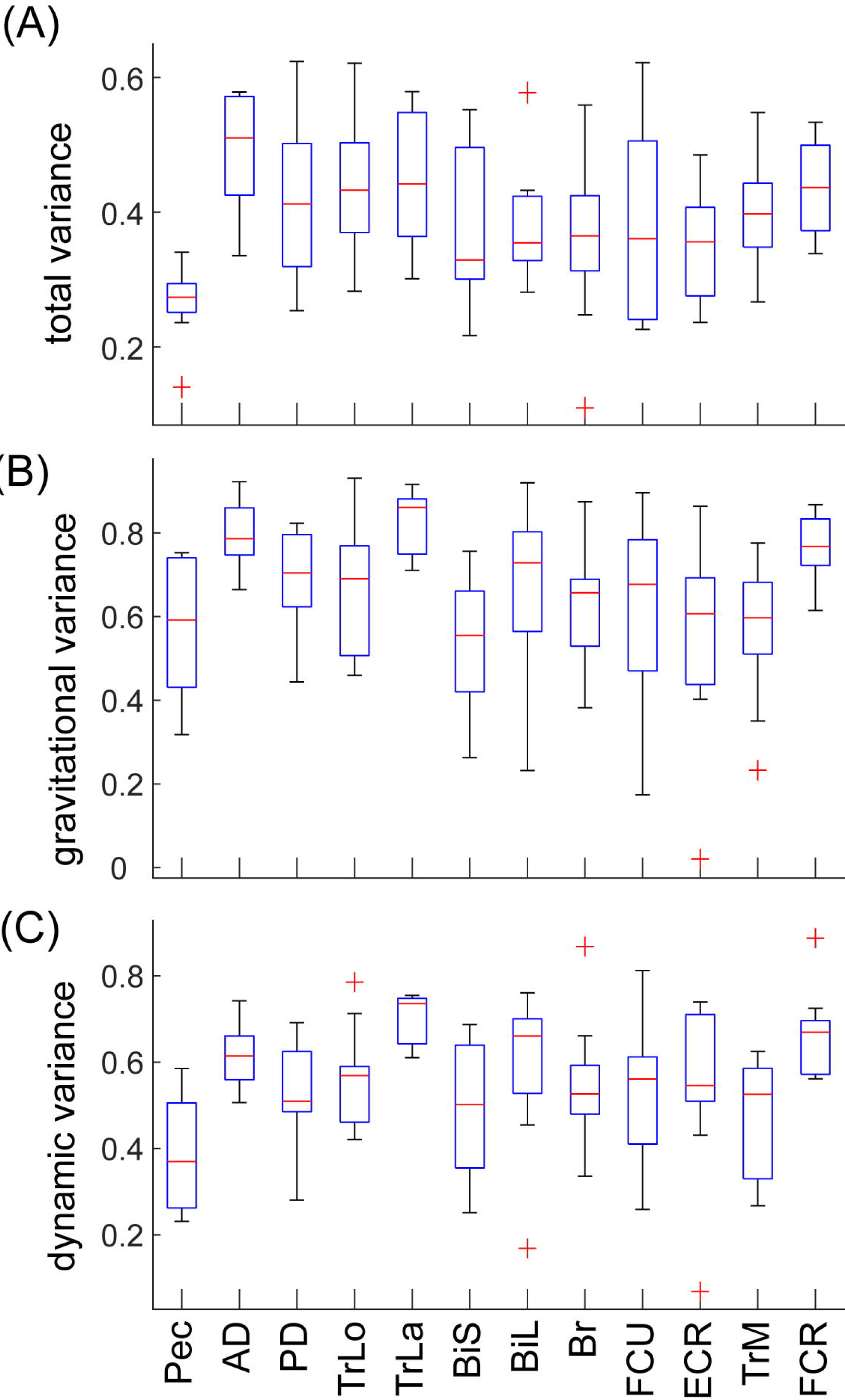
Variance accounted for by the individual components in EMG per muscle. Values are means across subjects and signals; boxes indicate ranges of 25 and 75 percentiles in data; error bars show standard deviations; red crosses indicate outliers. (A) Total variance accounted for by torque decomposition of EMG per muscle. (B) Relative variance accounted for by the gravitational principal component. (C) Relative variance accounted for by the dynamic principal component.

For movements in different directions, the muscle torques are accompanied by different amounts of relative contribution from MG and MD torques. We found that there were many instances, in which one or the other torque component tended to dominate the overall muscle torque, which is reflected in a distribution of r^2^ values along the maximal and minimal values (Fig. 6, blue). However, this was not the case for the distribution of r^2^ values for the contributions of torque principal components to EMG (Fig. 6, red). Surprisingly, the relative contributions of both torque principal components to EMG varied together in all muscles across all movement directions. This relationship was well fitted with a linear regression (p < 0.001 for all subjects), the slopes of these regressions ranged from 0.65 to 0.87 across subjects.

**Figure 6.**
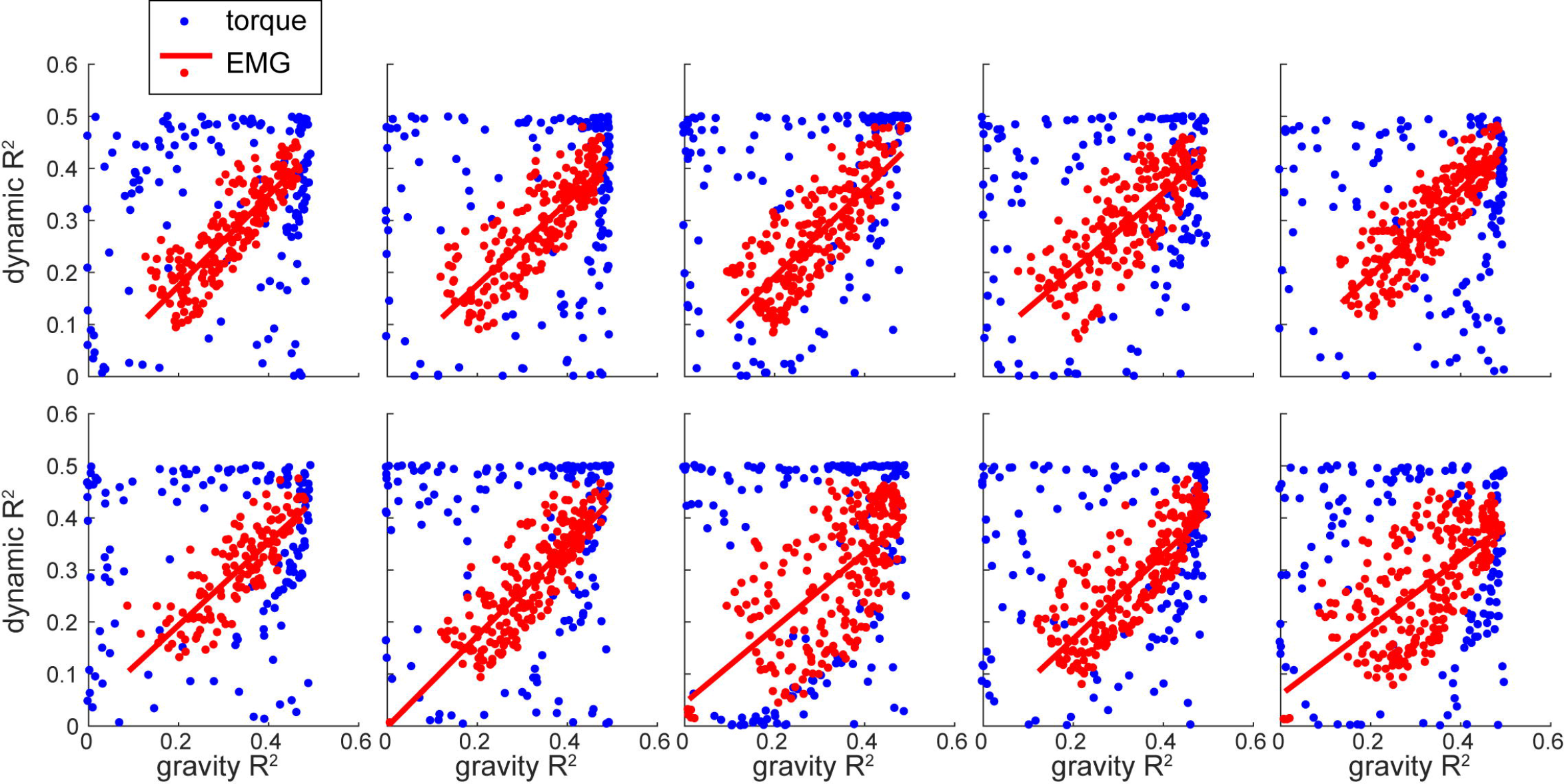
The relative contribution of individual components to EMG and muscle torques. Each plot shows data for a single subject. Each dot represents a value per signal (muscle or DOF) per movement.

## Discussion

The transformation from muscle activation to motion is non-linear and includes second order differential dynamics. This dynamics is often thought to be imbedded by the CNS either in the forms of internal models (Gomi and Kawato, 1997; Lackner and Dizio, 1994; Sabes, 2000; Shadmehr and Mussa Ivaldi, 1994; Wolpert and Kawato, 1998) or neural primitives (or synergies) (Bizzi et al., 1991; Giszter et al., 1993; Mussa Ivaldi, 1999; Mussa Ivaldi and Bizzi, 2000); for a review see (d’Avella and Lacquaniti, 2013). Here, we did not study the organization of the neural controller directly. Instead, we used the causal relationship between muscle contraction and motion to investigate a possible underlying relationship between limb dynamics and motor control mechanisms. Therefore, we will discuss our results within a context of a general control scheme that combines multiple views on the organization of the motor control system (Fig. 7). We have found that the gravitational and dynamic components of muscle torque explain on average 40% of EMG waveforms during goal-directed reaching movements, with more variance accounted for by the gravitational component than the dynamic component. This is further supported by the similarity between the first 2 principal components obtained from EMG and the first principal components obtained from gravitational and dynamic torque components separately. This is consistent with previous work showing that the first two principal components in EMG are related to static and phasic components of EMG waveforms (Flanders and Herrmann, 1992). Thus, the hypothetical neural commands originating in the supraspinal structures of the CNS and then combining with sensory feedback may form gravitational and dynamic components (Fig. 1). These commands could then underlie phasic and tonic components of EMG (d’Avella et al., 2008; Flanders, 1991). The supraspinal command compensating for gravity may constitute an anticipatory postural adjustment that accompanies movement that originates in the brainstem (Massion, 1992) or in the cerebellum (Sajdel-Sulkowska, 2013). Alternatively, or additionally, the gravitational command may be a spinal feedback response to changing gravitational load signaled by proprioceptors. The mechanism responsible for the feedback-driven compensation for gravity may be akin to positive force feedback during locomotion based on afferent feedback from Golgi tendon organs to maintain load bearing (Pearson and Collins, 1993; Prochazka et al., 1997). These components of neural commands, wherever they originate, are thought to optimally combine for motor control (Chhabra and Jacobs, 2006; Gaveau et al., 2014; 2016; Vu et al., 2016).

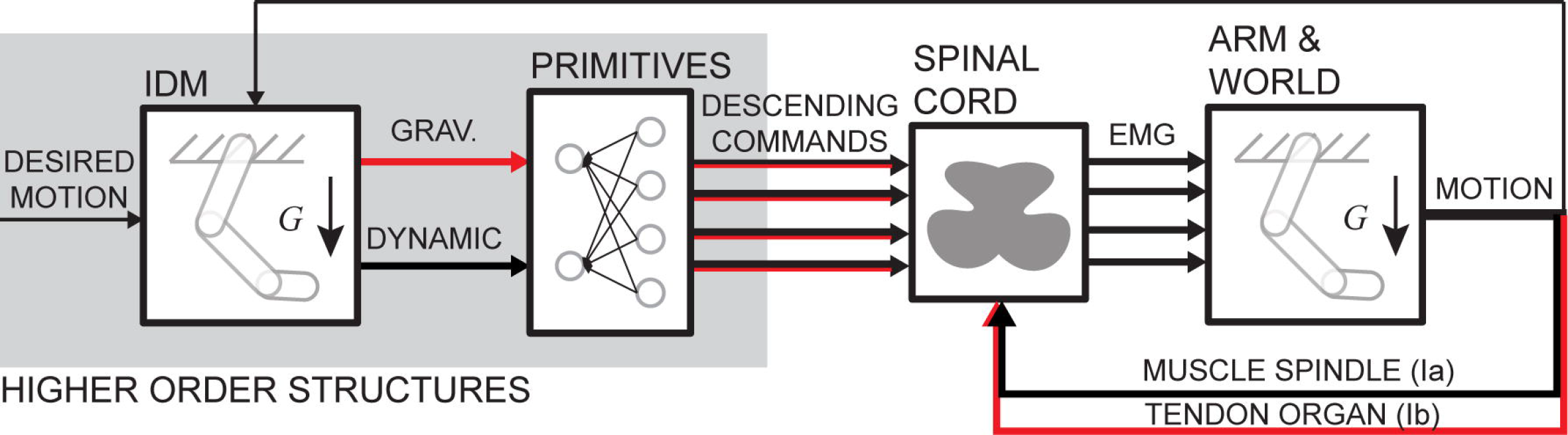

Our results indicate a linear relationship between the variance accounted for by the gravitational and dynamic torque components in EMG that was robust across subjects (Fig. 6). This relationship is not a direct reflection of the relative contribution of torque components to muscle torque. Instead, it may be a consequence of coupled torque waveforms across multiple DOFs and movement directions. In our data, this coupling is revealed by a single principal component that accounts for most of the variance in MD and MG torque waveforms. Similar single principal components were reported to capture most of the variance in kinematics and in dynamic torques during whole body reaching movements (Thomas et al., 2005). Although, our results show that the kinematic waveform in the Thomas et al. (2005) study may also be consistent with the gravitational torque waveform. Furthermore, the similarities in the waveforms of dynamic torques across DOFs and movements have been previously reported and interpreted as evidence of central planning (Gottlieb et al., 1997; Hollerbach and Flash, 1982; Thomas et al., 2005). Unexpectedly, our results may indicate that the coupling of the dynamic and gravitational torques across movements and DOFs may be reflected in muscle activity waveforms as nearly equal contribution of each of these components to EMG. This observation may suggest that the musculoskeletal structure, such as muscle moment arms around joints, muscle properties and composition of motor units, would then account for the scaling of muscle activity into appropriate moments (Gritsenko et al., 2016). However, large amount of variance was not accounted for by torque components in some muscles for some movements. Therefore, future studies will need to test this idea more directly.

The limitation of using joint torques to study the CNS is that they do not capture the effects of muscle co-contraction, which is an important control strategy used to alter joint stiffness or whole arm impedance (Damm and McIntyre, 2008; Darainy et al., 2004). However, our approach may offer some insight into this control strategy by using decomposition methods. Co-contraction implies a common signal across several antagonistic muscles. This signal is identifiable by PCA, which found 3 principal components that together accounted for more than 87% of variance in EMG. Our results indicate a close relationship between the first principal component of EMG and TG component of muscle torque (Fig. 3A). Furthermore, the contribution of gravitational torque component to EMG was not preferential to antigravity muscles, instead it was rather high across all muscles (Fig. 5). Similarly, the second principal component of EMG and TD component of muscle torque are similar (Fig. 3A) and the latter contributes to EMG across all muscles about equally (Fig. 5). This suggests that co-contraction may be part of the control strategy to compensate for gravitational and dynamic forces acting on the limb. Alternatively, co-contraction may be captured by the third principal component of EMG, which has an even more phasic waveform than the second principal component. Future studies may obtain further insight into this issue by using our method to account for the contribution of muscle torques to EMG.

## Conclusions

In conclusion, our results have shown that gravitational and dynamic components of muscle torque represent significant amount of variance in muscle activity. This suggests that these forces may underlie phasic and tonic components of muscle activity.

## Acknowledgments

The authors wish to thank Dr. Sergiy Yakovenko for his contribution to the discussion of analysis in this study and Dr. Robert L. Goodman and Dr. Amy J. Bastian for their critical review of this manuscript.

## Financial Disclosure

This research was sponsored by NIH/NIGMS U54GM104942 (EO) providing student fellowship, NIH P20GM109098 providing salary support (VG, BP) and supplies, NIH P30GM103503 providing equipment support. The content is solely the responsibility of the authors and does not necessarily represent the official views of the NIH.

